# Face-selective units in human ventral temporal cortex reactivate during free recall

**DOI:** 10.1101/487686

**Authors:** Simon Khuvis, Erin M. Yeagle, Yitzhak Norman, Shany Grossman, Rafael Malach, Ashesh D. Mehta

## Abstract

Properties of face-responsive individual neurons in the human ventral temporal cortex (VTC) have yet to be studied, and their role in conscious perception remains unknown. To explore this, we implanted microelectrodes into the VTCs of eight human subjects undergoing invasive epilepsy monitoring. Most (26 of 33) category-selective units showed specificity for face stimuli, with a range of response profiles. Different face exemplars evoked consistent and discriminable responses in the population of units sampled. During a free recall task, face-selective units selectively reactivated in the absence of visual stimulation during the 2-second window prior to face recall events. Furthermore, the identity of the recalled face could be predicted by comparing activity preceding recall events to activity evoked by visual stimulation.

## Introduction

Facial recognition is an essential adaptive social function in primates, facilitated by the extensive development of specialized visual areas in the brain’s ventral temporal cortex (VTC). Information processing in this region must meet social demands to recognize, classify and uniquely identify a multitude of faces. First described in monkeys, studies of single neuron firing in response to visual features of faces have uncovered key “bottom-up” mechanisms of the feature space that drive neuronal firing^1–3^. VTC neurons exhibit precise facial feature sensitivity, supporting their role in the discrimination of individual face exemplars. However, their role in non-sensory, extraretinal processing, as would occur during imagery and recall, is difficult to determine from monkeys, who cannot qualitatively express their experiences. In humans, neuroimaging has defined a critical node in face processing within the VTC: the fusiform face area (FFA)^4^. Functional MRI (fMRI) studies support both a sensory (“bottom-up”) as well as a cognitive (“top-down”) role for the FFA, which is activated not only when subjects view faces, but also when they expect to see a face^5,6^, perform imagery tasks involving faces^7,8^, and hold face representations in working memory^9^. Activation of category-selective regions of VTC can predict recall of items in that category^10,11^. In addition to fMRI studies, magnetoencephalography^12^ and direct electrocorticographic recordings^13^ in humans show fusiform responses emerging early (~100 ms) after face presentation, suggest a sensory role for this region. While these studies have provided important insights, they lack the spatiotemporal resolution needed to uncover how bottom-up, feature-based, sensory processing relates to top-down processes at the level of single human neurons. Thus, the exquisite neuronal selectivity for facial features revealed by monkey neurophysiology, the relatively early sensitivity revealed by human field potential recordings and the fMRI evidence for top-down control of the FFA have yet to be integrated.

Clinical macroelectrodes with microwires provide an opportunity to investigate this question by studying neuronal spiking in patients undergoing chronic recordings. Such *in vivo* human data have provided insights into a number of physiological and pathological processes, most notably in the medial temporal lobe^14^. Face-responsive units have been reported in the human VTC^15^, but their single and population spiking activity has yet to be explored in detail. In experiments described here, we recorded from microwires in the VTCs of human subjects as they viewed face stimuli and, later, recalled them in an episodic free recall task, allowing us to examine human sensory and higher-order cognitive processes at the single-unit level. Recorded units showed a diversity of response patterns while maintaining strong category specificity. We show that population responses to the presentation of different face exemplars can be robustly discriminated, and that these responses are reinstated when subjects recall and visualize previously-presented face images in the absence of sensory input. Our results support models of memory in which single-neuronal substrates of sensory processing are reactivated in a top-down fashion during recall^16^.

## Results

### Face-Selective Units Found in Ventral Temporal Cortex

Eight subjects (four females, see Table 1) with partial epilepsy, undergoing diagnostic intracranial EEG (iEEG) monitoring of VTC were implanted with micro-macro depth electrodes^17^ (Ad-Tech Medical, Oak Creek, WI). All subjects gave informed consent under IRB guidelines, and electrode targeting was guided primarily by clinical needs. Typically, this was to help determine the posterior margin of resection in patients with putative temporal lobe epilepsy in an effort to preserve the VTC and avoid deficits in object identification. Subjects performed a 1-back task requiring the identification of face, body part, house, tool and pattern exemplars (see Fig. S2) while undergoing functional MRI before surgery. Electrode positions were localized post-operatively by superimposing MRI and computed tomography scans^18^; an example is shown in Fig. 1A (full data, Fig. S1). None of the VTC areas sampled with microelectrodes were involved in seizure onset, as determined by epileptologist evaluation. Subjects repeated the 1-back task (Fig. S2) while microelectrode data were recorded at 30 kHz. Sixty-three visually-responsive single and multi-units were recorded collectively from all eight subjects of 124 total units (see Table 1 for single-subject data). Of these, 33 were category-selective, with the vast majority (26, recorded from five subjects, 41% of visually-responsive units) selective for faces (see Fig. S3). The average time-course of all visually responsive units across subjects showed a marked preference for faces (Fig. 1B), with onset latencies of face-selective units distributed between 100 and 220 ms after presentation of face stimuli (Fig. 1C). Visual responsiveness was strongly correlated with face selectivity (Fig. S4, *p* < 0.001, Spearman’s ρ, *N* = 120 units in subjects who performed the standard version of the task).

**Fig. 1.**
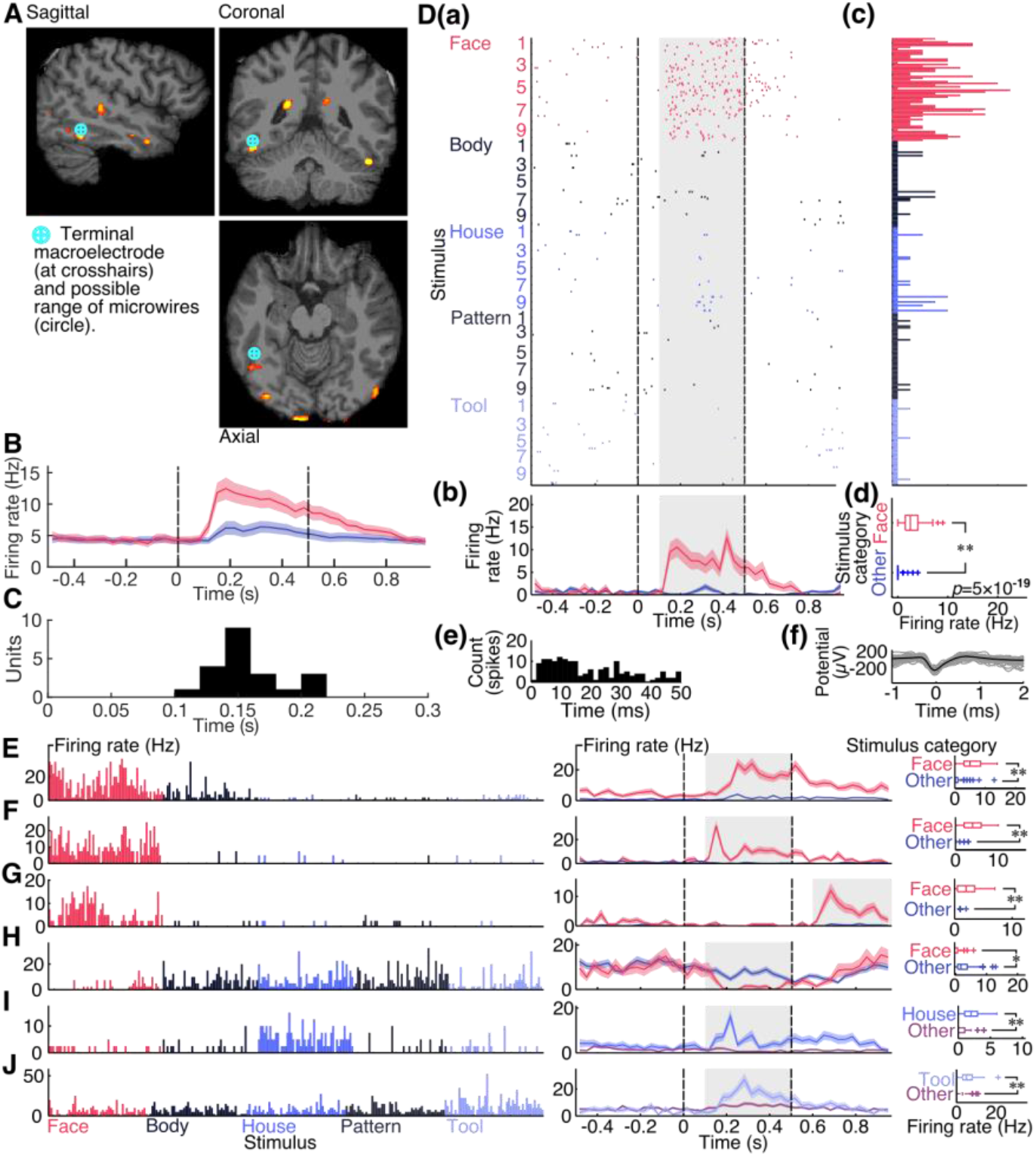
Diverse category-selective units found in the ventral temporal cortex. (A) Example electrode location (Subject 8) over functional MRI face>house activation contrast map (in red). (B) Grand mean peristimulus-time histogram. Red: faces, blue: non-face objects. Colored shaded areas represent mean ± standard error of the mean (SEM) at each timepoint. Dashed lines represent image onset at time 0 and offset, respectively. (C) Onset latency of face-selective units. (D) Representative face-selective unit from right FFA in Subject 8. (D(a)) Raster plot of responses with each row representing a single trial, and dashed lines representing onset and offset respectively. (D(b)) Mean peristimulus time histogram. Red: faces, blue: non-face objects. Colored shaded areas represent mean ± SEM at each timepoint. (D(c)) Mean firing rate per trial. Average over gray shaded area in raster plot, 0.1 to 0.5 s after presentation. (D(d)) Distribution of responses to faces and non-face objects. Responses to faces are significantly stronger. (D(e)) Inter-spike interval distribution. (D(f)) Spike waveforms. Mean spike waveform in black. (E-J) Peristimulus time histograms and grand mean peristimulus time histograms for units with (E) longer response latency, (F) transient peaked response, (G) offset response, (H) suppression to faces, (I) house selectivity, (J) tool selectivity. *: *p*<0.05; **: *p*<0.001, rank-sum test, Bonferroni-Holm correction.

**Table 1.** Demographic and recording information for all participating subjects. Sub: subject, Hand: handedness, Lang: primary language, FSIQ: full-scale IQ, as measured by the Wechsler Adult Intelligence Scale, Fourth Edition, Pearson Clinical (London), Faces: Face (Supplemental) scaled score from Pearson Clinical (London) Social Cognition test – a test that requires intact facial recognition, memory and concentration (mean of 10, standard deviation of 3), Imp: implant number, Lat: laterality of electrode, Vis: visually-responsive units, F: face-selective units, B: body-part-selective units, T: tool-selective units, H: house-selective units, P: pattern-selective units (the sole pattern selective unit shows a weaker response to patterns than all other stimulus classes), S: scene- (place) selective units. Eng: English, Span: Spanish. NT: not tested. R: right, L: left. M: male, F: female.

| Sub | Sex | Age | Hand | Lang | FSIQ | Faces | Imp | Lat | 1-Back |  |  |  |  |  | Free Recall |  |  |
| --- | --- | --- | --- | --- | --- | --- | --- | --- | --- | --- | --- | --- | --- | --- | --- | --- | --- |
|  |  |  |  |  |  |  |  |  | Vis | F | B | T | H | P | Vis | F | S |
| 1 | F | 32 | R | Eng | 67 | 4 | 1 | L | 2 | 2 | 0 | 0 | 0 | 0 |  |  |  |
| 2 | M | 41 | L | Span | N/A | N/A | 1 | R | 7 | 0 | 0 | 0 | 0 | 0 | 14 | 0 | 2 |
|  |  |  |  |  |  |  | 2 | L | 4 | 1 | 1 | 0 | 0 | 0 | 10 | 5 | 0 |
| 3 | F | 55 | R | Eng | 46 | NT | 1 | R | 8 | 8 | 0 | 0 | 0 | 0 | 12 | 7 | 0 |
|  |  |  |  |  |  |  |  | L | 1 | 0 | 0 | 0 | 0 | 0 | 2 | 0 | 0 |
| 4 | F | 37 | R | Eng | 74 | 2 | 1 | L | 5 | 0 | 0 | 1 | 0 | 0 |  |  |  |
| 5 | M | 20 | R | Span | N/A | N/A | 1 | R | 8 | 0 | 0 | 0 | 2 | 1 |  |  |  |
| 6 | F | 32 | R | Eng | 88 | 10 | 1 | R | 13 | 3 | 0 | 0 | 2 | 0 | 8 | 0 | 1 |
| 7 | M | 43 | R | Eng | 91 | 12 | 1 | R | 2 | 0 | 0 | 0 | 0 | 0 |  |  |  |
|  |  |  |  |  |  |  |  | L | 1 | 0 | 0 | 0 | 0 | 0 |  |  |  |
| 8 | M | 20 | R | Eng | 77 | 11 | 1 | R | 12 | 12 | 0 | 0 | 0 | 0 | 16 | 16 | 0 |

Fig. 1D shows a typical face-selective unit, with a vigorous response to face image presentation and return to baseline firing rate after the image disappeared. However, we also observed a surprising diversity of face-selective response patterns, including units with response persistence (Fig. 1E, Fig. S5) and units whose activity showed sharp transient peaks after face presentation (Fig. 1F, Fig. S6). One unit showed a strong face-selective offset response (Fig. 1G, Fig. S7). Several units showed selective suppression to face presentation (Fig. 1H, Fig. S8), a finding consistent with single-unit recordings from inferotemporal cortex in non-human primates^19,20^.

We also recorded units with selectivity for non-face categories – tools and houses. With the exception of two house-selective units in one subject, from whom only weakly face-selective units were recorded, all house- and tool-selective units came from subjects whose recording sites yielded no face- or body-selective units. This is consistent with the reported segregation of domains dedicated to processing animate and inanimate objects^21,22^. After face-selective units, house-selective units were the most common, with four (6% of visually-responsive units). Fig. 1I shows one such unit (also see Fig. S9). The sole tool-selective unit is shown in Fig. 1J (also see Fig. S10).

### Exemplar Decoding of Faces in Ventral Temporal Cortex Ensembles

Next, we sought to corroborate previous reports suggesting that individual faces (face exemplars) had unique representations in the FFA^23–25^, in human VTC more broadly^26^, and, at the single-unit level, in macaque face patches^1–3^. Responses to single presentations (trials) of exemplars from all visually-responsive units across all subjects were concatenated into a single log-transformed pseudo-population vector. Multi-dimensional scaling (MDS), a linear technique for dimensionality reduction, was then applied to visualize the relationships among trials of different exemplars in a common space. For example, responses from all trials to three face and house exemplars (Face 1–3 and House 1–3 in the stimulus set, chosen *ex ante*, for illustration) are presented in Fig. 2B. As expected, we observed a clear segregation of faces and houses in the representational space. We then applied MDS to the three face exemplars (Fig. 2C) and the three house exemplars (Fig. 2D), alone, and found that trials of individual face exemplars appeared to be linearly separable, while trials of house exemplars were not.

**Fig. 2.**
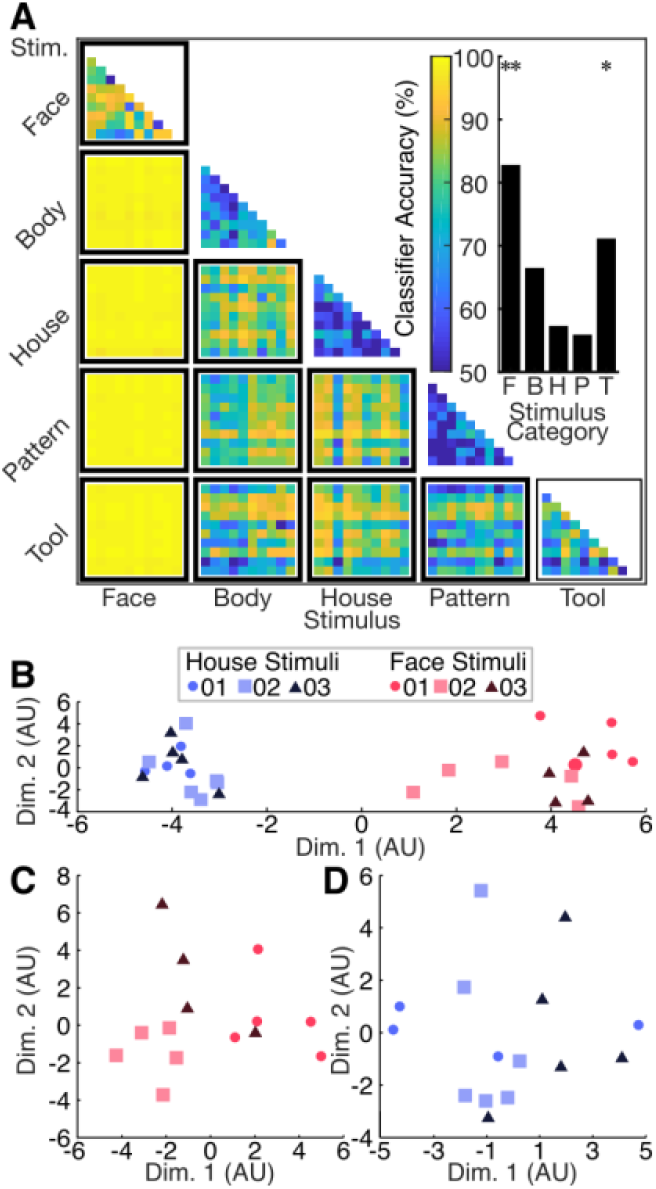
Exemplar decoding of face stimuli. (A) Binary classifier performance. Cross-validation accuracy of a set of classifiers tested and trained on each pair of exemplars. Discriminability and classifier accuracy of any exemplar pair represented by the color at that location in the matrix, with greater accuracy corresponding to more dissimilar population responses. Inset: Mean decoding accuracies for each within-category block (classifier accuracy discriminating exemplars of the same category). */thin outline: *p*<0.05; **/thick outline: *p*<0.01, bootstrap test, Bonferroni correction. (B) Multidimensional scaling: population responses to trials of first three face and house exemplars. (C) Multidimensional scaling: population responses to trials of first three face exemplars. (D) Multidimensional scaling: population responses to trials of first three house exemplars.

To quantify this, we plotted transformed pseudo-population response vectors from each pair of exemplars presented and used a simple linear discriminant to separate them. We then performed leave-one-out cross-validation as a measure of the discriminability of each of these exemplar pairs (Fig. 2A). Classifier accuracy was very high for exemplar pairs from different stimulus categories, especially faces *vs*. non-face objects, showing that responses to these categories are distinct (*p* < 0.0001, bootstrap test, Bonferroni correction at *N* = 15 category pairs). While it is unsurprising that faces can be distinguished from non-faces, given the vastly different magnitudes of observed unit responses, the classifier was also able to discriminate between non-face categories, for example, tools and houses. Confirming the previous studies, we show robust exemplar selectivity, evidenced by strong classifier performance in discriminating face exemplar pairs (83%, *p* = 0.0001, bootstrap test, Bonferroni correction at *N* = 5 categories). We also found some weaker tool exemplar decoding (73%, *p* = 0.008, bootstrap test, Bonferroni correction at *N* = 5 categories), but no within-category exemplar decoding for other categories (bodies: 66%, *p* = 0.03; houses: 57%, *p* = 0.2; patterns: 56%, *p* = 0.3).

### Free-Recall- and Imagery-Evoked Reactivation of Face Representations

Next, we tested whether face representations in VTC could be activated endogenously in the absence of external visual input. Specifically, we wanted to learn whether VTC units that responded selectively to viewing of face images would respond in a similar way when subjects recalled or imagined those same face images. To that end, four subjects also performed an episodic free recall task^11^, in which they were shown and asked to remember full-color photographs of famous faces and scenes. After performing a short interference task and putting a blindfold on, the subjects were asked to freely recall as many pictures as possible, focusing on one category (faces/places) at a time. The subjects were instructed to visualize and describe each picture they recalled in as much detail as possible, emphasizing unique colors, facial expressions, lighting, perspective, *etc*. Face-selective units were recoded from 3/4 of this subset of subjects, and small numbers of place-selective units were recorded from 2/4 (also see Fig. 4A).

Guided by prior fMRI results, we first sought to identify whether face-selective units were reactivated during visual imagery of faces^7,8^. We examined mean firing rates of VTC units within two seconds of the verbal recall event, reasoning that activity associated with visualization most likely occurred in this window. Units’ firing rates during face presentation (over pre-stimulus baseline, see *Methods*), and in the 4-second interval centered at onset of the face recall utterance (over whole-experiment baseline), were well correlated (*p* = 0.008, Spearman’s ρ = 0.33, *N* = 62 units, Fig. 3B), as were their preferences for face stimuli during presentation and recall (*p* = 0.003, Spearman’s ρ = 0.37, *N* = 62 units, Fig. 3C). This correlation persisted when only strongly visually-responsive units (see *Methods*) were included (*p* = 0.002, Spearman’s ρ = 0.55, *N* = 29 units). To further examine this content-selective relationship, and to investigate the precise temporal dynamics of this recall-triggered activity, we computed the average baseline-corrected activity for each presentation trial (Fig. 3D) and each recall event (Fig. 3E) for all face-selective units in each implant, and compared the face to place stimuli. Activity in face-selective units was significantly greater around face recall events than around place recall events (unpaired *t*-test, *p* = 0.02). Mean activity in face-selective units began increasing around 2 seconds before onset of a face recall utterance, peaked, and returned to near baseline as the subject began to speak. This temporal relationship is consistent with fMRI^10^ results that demonstrate activation of face-selective brain areas, including the FFA, during recall, and iEEG results^11^ showing an increase in the high-frequency broadband signal in category-selective VTC in the seconds leading up to recall of an item in that category.

**Fig. 3.**
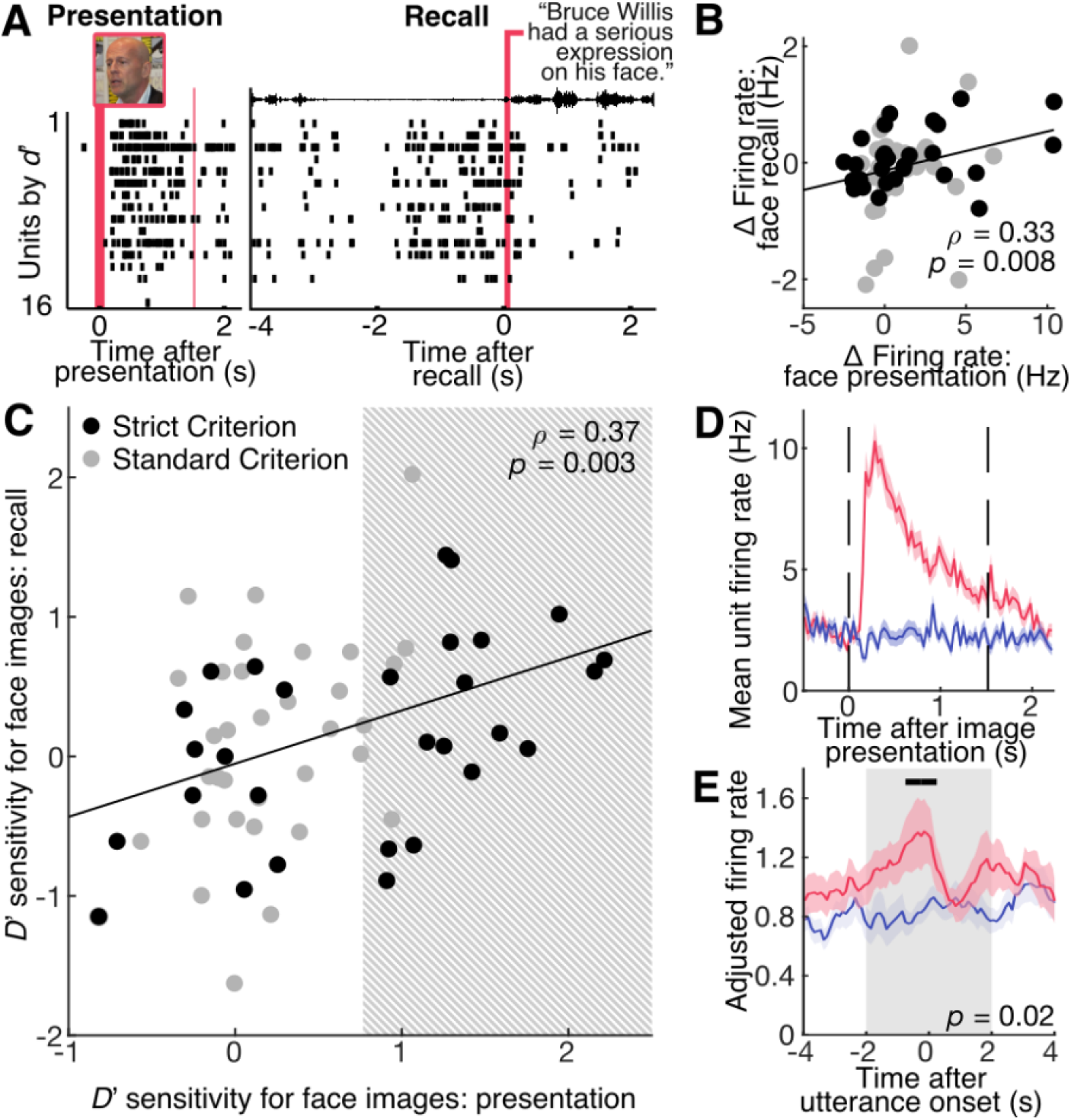
Face-selective units reactivate during recall. (A) Raster plot from presentation and recall example face stimulus from Subject 8. Image presentation of actor Bruce Willis^38^ at time 0 on left, and vocalization onset (black envelope trace at top) on the right. (B) Responses to faces during presentation and before recall. Black: units passing stricter visual responsiveness criterion. (C) Selectivity for faces during presentation and recall. Trend line fitted to all units. (D) Mean peri-stimulus time histogram for face trials. Face selectivity defined by gray area in (C). Responses to faces in red and places in blue. Colored areas: mean ± standard error of the mean at each timepoint across subjects and trials (*N*_1_ = *N*_2_ = 140 place and face presentation trials). Dashed lines represent image onset at time 0 and offset. (E) Mean peri-recall time histogram. Significant difference between face and place activity (gray box, −2 to 2 seconds, two-tailed unpaired *t*-test, *p* = 0.02, *t*(48) = 2.4, *N*_1_ = 24 face recall events, *N*_2_ = 26 place recall events). Black bar: two-tailed unpaired *t*-tests, *p* < 0.05, *t*(48) > 2.01, uncorrected.

Next, we sought to determine whether neural ensembles from subjects with selectivity for individual face exemplars would demonstrate similar selectivity during recall. Figure 3A (and Fig. S11) shows single-trial raster plots for face-selective units during stimulation and recall. Patterns of activity before recall were strikingly similar to those observed during initial face presentation, suggesting that it might be possible to predict the identity of the face to be recalled by matching activity of these units before a recall event to their activity during presentation. To examine this possibility, we first identified all subjects whose recorded units showed above-chance exemplar decoding during presentation for each category. This reproduced the findings from the 1-back task (Fig. 2) for each subject (Fig. 4B), with the exception that each pseudopopulation vector was first normalized to account for the anticipated potential differences in magnitude of activation between presentation and recall periods (see *Methods*, Fig. S12). Mean leave-one-out cross-validation accuracy was calculated for classification among exemplars of the same category (“within-category”) and all exemplars (“cross-category”). All subjects exhibiting face selectivity (Subjects 2, 3 and 8) showed above-chance cross-category classification accuracy (Fig. 4B, bootstrap test, *p* < 0.05, Bonferroni-Holm correction at *N* = 4 subjects, Table S1 for exact *p* values). Subjects 3 and 8 also showed above-chance within-category face classification accuracy. Subject 6 had no face-selective units (Fig. 4A), and predictably, no exemplar decoding.

**Fig. 4.**
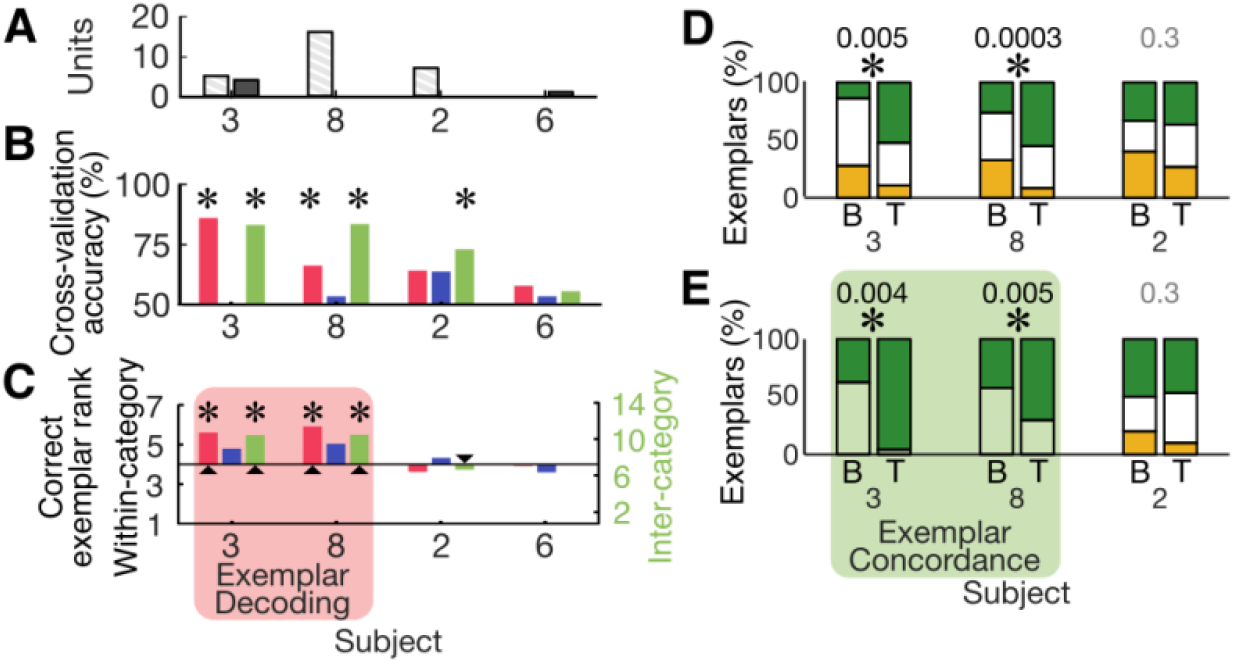
Exemplar decoding of recalled stimuli. (A) Number of face- (light, hashed bars) and place- (dark bars) selective units recorded in each subject in whom the experiment was performed. (B) Presentation exemplar decoding in recall task data. Red bars: faces; blue bars: places; green bars: cross-category decoding. *:*p* < 0.05, bootstrap test, Bonferroni-Holm correction. (C) Recall exemplar decoding. Red bars: faces, left axis; blue bars: places, left axis; green bars: cross-category decoding, right axis; arrowheads: subject-categories with presentation exemplar decoding; red shaded subjects: subjects with face presentation exemplar decoding (in whom face recall exemplar decoding is expected). *: *p* < 0.05, bootstrap test, Bonferroni-Holm correction in subject-categories with significant presentation exemplar decoding. (D) Mean concordance between presentation response and recall activity terciles among all exemplars, across face-selective units. B: exemplars in bottom tercile of recall activity; T: top tercile. Green fraction: exemplars in top tercile of presentation response; yellow fraction: bottom tercile. *: *p* < 0.05, bootstrap test, Bonferroni-Holm correction. Numbers above bars: uncorrected *p* value. (E) Mean concordance between presentation response and recall activity terciles among face exemplars, across face-selective units. Green shaded subjects: subjects showing concordance among all exemplars (in whom face concordance is expected); otherwise, labeling as above.

For categories that showed above chance decoding accuracy (Fig. 4C, arrowheads), we then tested whether the same classifiers, trained on the presentation data, would be able to decode exemplar identity based on activation prior to a recall event (*i.e*. cross-classification analysis). Both Subjects 3 and 8 (the two who showed exemplar decoding during presentation) showed significant exemplar decoding during recall for faces and across categories, both in combination (*p* = 0.03, cross-category and *p* = 0.0017 for faces, only, bootstrap test, Fisher’s method of combined probabilities) and individually (*p* < 0.05, bootstrap test, Bonferroni-Holm correction at *N* = 2 and 3 tested subjects for faces and cross-category respectively, Table S2 for exact *p* values). Predictably, the excluded subject that did not demonstrate exemplar decoding during stimulation (Subject 2) also did not demonstrate it during recall.

To further confirm that face-selective units showed a consistent exemplar tuning between presentation and recall, we examined the proportion of exemplars in the top and bottom terciles of recall activity amplitude (measured using mean firing rates 2–0 seconds before recall utterance) for each face-selective unit that also evoked presentation responses in the top and bottom terciles, respectively, for that unit. Units with a large proportion of exemplars in the top and bottom terciles during recall that were also ranked in those same terciles during presentation have a strong concordance between presentation and recall tuning. As with the cross-classification analysis, both Subjects 8 and 3 (the two who showed exemplar decoding during presentation) showed above-chance concordance among face-selective units, among exemplars of both categories (“cross-category concordance”, Fig. 4D, *p* < 0.05, bootstrap test, Bonferroni-Holm correction at *N* = 3 subjects). We then tested for concordance among only face exemplars. Again, the same subjects (8 and 3) showed significant concordance between presentation and recall among face exemplars (Fig. 4E, *p* < 0.05, bootstrap test, Bonferroni-Holm correction at *N* = 2 subjects with significant cross-category concordance).

## Discussion

We present the first comprehensive study of single-unit firing properties of the human VTC. Units in the vicinity of the FFA show a diverse range of responses to visual stimulation, including transient, sustained, suppression and offset responses. While suppression from baseline firing has been documented in area IT in non-human primates^20,27^, we are not aware of any reports of offset responses in higher visual areas of any animal model. In addition to face-selective units, we also recorded several units responsive to houses and patterns in three subjects. In all but one of these (Subject 6), no face- or body-selective units were observed. In Subject 6, only weakly-face selective units were recorded. This is consistent with the previously-reported segregation of animate and inanimate object representations in VTC.^21,22^

Next, we demonstrated that representations of individual face exemplars were invariant across trials. To do this, we used multidimensional scaling to show that repeated presentations of the same exemplar evoke similar responses in the neuronal ensemble, and that a classifier can consistently identify exemplars within a category at a rate above chance. Face exemplars were strongly discriminable, and tools also showed weaker, but above-chance, discriminability. The tool exemplar decoding may have been a product of the single tool-selective unit in the pseudo-population data set. Alternatively, the vastly different shapes of the tool stimuli may have evoked subtle differences in the responses in units tuned to other stimuli. As has been argued by Kanwisher and colleagues^28,29^, simply decoding stimuli from the activity of a neuronal ensemble is not, in itself, proof that that ensemble is dedicated to representations of such stimuli.

To circumvent this experimental limitation, subjects were asked to report on, and visualize, freely-recalled images, and we compared responses in the purported face area to those evoked when subjects initially viewed those same images. This experiment helps to evaluate the claim that face representations are native to VTC, as it is free of the confounding influence of visual input. In general, units’ face selectivity around recall events was correlated to their selectivity around presentation, and a mean of the activity of face-selective units across face and place recall events (Fig. 3F) showed that this was driven by a face-specific transient increase in face unit activity in the 2 s preceding recall (note that this analysis minimizes the influence of clustering decisions by first collapsing across all face-selective units in each subject). It was these 2 seconds of data that were subjected to the recall exemplar decoding analysis.

Two subjects: 8 and 3, showed above-chance exemplar decoding during presentation (*i.e*. a classifier was able to discriminate population codes for specific face exemplars when these subjects viewed those exemplars). Subjects 2 and 6 also performed the experiment, but the recordings from Subject 6 yielded no face-selective units during the free recall task, suggesting the electrode was outside of a core face processing region. Subject 2 also showed no exemplar decoding during presentation – this may also have been because activity in the face-selective units in this subject was merely correlated with face processing in general, and was not causally involved in face representation. This would be expected if the electrode lay in any of the broader VTC brain areas activated by face presentation on fMRI, but not within a core face-representation region (*e.g*., purportedly, the FFA). The fact that this very same electrode also recorded a body-selective unit in the 1-back task performed at an earlier time (see Table 1) suggests that units from this area belong to an adjacent network activated during viewing of animate objects generally, rather than faces in particular. The free recall task did not include presentation of non-face animate objects, so it would be impossible to tell if these units were indeed specific for faces. In contrast, electrodes in Subjects 3 and 8 recorded only face-selective units in the 1-back task.

Regardless of the reason, Subjects 2 and 6 act as negative controls – if no exemplar decoding was found during presentation, none would be expected during recall. In contrast, subjects with exemplar decoding during presentation should exhibit such decoding during recall. This is the pattern we observed. Furthermore, a classifier trained to recognize exemplars based on activity during presentation could successfully decode exemplars based on activity during recall, indicating that recall activity was roughly a reinstatement of presentation activity. This was true both at an ensemble level and at the level of individual units, whose tuning preferences for different exemplars were conserved between presentation and recall (see Fig 4E).

Our findings illustrate that face-selective units are recruited in category-specific recall and/or imagery. Though we do not prove causality in this experiment, our work suggests that the same brain areas are responsible for face processing during viewing and in face recall/imagery. More broadly, it aligns with the hypothesis that subjective percepts are supported by common category-specific brain areas, whether they are invoked by external stimulation or by internal reactivation.

In summary, we demonstrate a detailed description of single unit properties from the human VTC responsive to face and non-face stimuli during viewing and during an imagery task.

Extending prior observations from nonhuman primates, we report a diverse range of highly face-selective units within human VTC; that those units form a population code by which individual face exemplars can be discriminated; and that reactivation of the patterns forming that code occurs not only during face perception, but also during face imagination and recall. In line with prior neuroimaging work supporting a role of the VTC in conscious perception, we demonstrate selective activation during recall at the single neuron level. These findings support the role of the VTC, and FFA specifically, as a critical substrate of conscious face representation, one used not only to identify and discriminate faces in the environment, but also to support face representations generated internally. Our research adds to a large and growing body of literature supporting a role for higher-order sensory areas in subserving working memory, imagery and other cognitive processes that engage the same neuronal substrates as bottom-up sensory processes.

## Methods

### Subject Recruitment

Study protocols were approved by the Institutional Review Board of the Feinstein Institute for Medical Research. Eight subjects (4 females) scheduled to undergo intracranial electroencephalography (iEEG) of the inferior occipitotemporal cortex, including either the right or left FFA, for diagnosis of focal epilepsy were recruited. The determination of which brain areas to monitor for each subject was made on strictly clinical grounds. One subject (Subject 2) was implanted twice: first on the right and then on the left. He performed the 1-back task twice, and three runs of the free recall task (typically there are two runs per subject).

Six subjects had electrodes on the right, 5 subjects had electrodes on the left (3 bilateral). Subject demographic and surgical data and recording yields are presented in Table 1.

### Electrode Placement

Subjects 3–8 performed the same 1-back task during an fMRI scan as that they did during intracranial single unit recordings (see Fig. S2, and *Tasks* for task details). Focal areas with the greatest Face>House contrast within the fusiform gyrus, and consistent with the expected anatomy, were identified as the FFA.

In the recruited subjects, monitoring of brain areas including the FFA at least unilaterally was clinically indicated for localization of epilepsy and mapping of functional areas. Precise electrode trajectories are typically underdefined by clinical circumstances, and when a range of *a priori* equally-efficacious and safe alternatives included a trajectory leading to either the right or left FFA, then this brain area was targeted either by (1) anatomical approximation based on pre-operative structural MRI in the Radionics electrode trajectory planning software (Integra LifeSciences, Plainsboro, NJ) by a neurosurgeon experienced in functional neuroanatomy (A.D.M.), based on landmarks known to define the FFA^22^, or by (2) at least two investigators (A.D.M. and S.K.) comparing the relative anatomy of the fMRI-defined FFA to the anatomy shown in the pre-operative MRI and superimposing the Face>House contrast map from the fMRI over the pre-operative MRI in the StealthStation trajectory planning software (Medtronic, Dublin, Ireland). The end of each macro electrode was planned to stop 4 mm short of the centroid of the targeted area so that the microwires, when deployed, would terminate in the region of interest.

Ad-Tech Medical (Oak Creek, WI) micro-macro depth electrodes were used to simultaneously obtain clinical EEG data as well as record single-unit activity with minimal additional surgical risk. Micro-electrodes were trimmed to 4 mm beyond the end of the macro canula prior to insertion. One micro-macro electrode was implanted in each targeted FFA, each with eight signal and one reference microwires.

### MRI Co-Localization

Six out of the eight subjects underwent a functional MRI scan in addition to the pre-implant anatomical image acquisition. Blood-oxygenation-level dependent (BOLD) contrast was obtained using a T2* sensitive echo planar imaging (EPI) sequence: TR 2000 ms; TE 30 ms; flip angle 77°; field of view 192 mm; voxel size 2.1×2.1×2.1 mm; 34 slices). During the functional scan, subjects performed the same 1-back task as performed during iEEG recordings (see Fig. S2 and *Tasks*).

FMRI data analyses were carried out using FSL’s FEAT^30^, version 6.00 (www.fmrib.ox.ac.uk/fsl). Preprocessing steps included discarding first two acquired volumes; motion correction using MCFLIRT, with the first volume as a reference; slice-timing correction of voxels time series using sinc interpolation; high-pass temporal filtering with a cutoff of 0.0083 Hz (maximal cycle of 120 s); and spatial smoothing of each acquired volume using a Gaussian kernel with full width at half maximum (FWHM) of 2mm.

A general linear model was fit to each subject’s run, with blocks from the five different visual categories modeled by five independent box car predictors convolved with a double gamma hemodynamic response function model. Motion correction time series, their first derivatives and their squared time series were included as additional confound predictors (18 predictors in total, 6 original motion series ×3). Each predictor and voxel’s time series was demeaned prior to computing the general linear model to account for baseline activity level. For each condition and contrast, we obtained a whole brain *t*-value map, and aligned the map to the high-resolution anatomical image of the subject using FLIRT.

Electrodes were localized using the iELVis toolbox^18^. A post-implant CT for each subject was registered to the pre-implant MRI *via* the post-implant MRI, using FSL’s FLIRT^31–33^. Following coregistration, electrode artifacts were identified manually in BioImageSuite^34^.

For visualization, functional data were coregistered to the subject’s brainmask volume (generated *via* FreeSurfer’s `recon_all`) using 3dAllineate (AFNI toolbox – afni.nimh.nih.gov)^35^ and thresholded (minimum *t*-value = 1.5–3; maximum *t*-value = 6, cluster threshold = 100 voxels). The contrast used was faces vs. houses, except for one subject (Subject 4) for whom faces vs. patterns yielded a clearer map. Macroelectrode coordinates were superimposed on the contrast map using BioImageSuite^34^.

### Data Acquisition

Neural data were acquired at 30 kHz on a Blackrock Microsystems (Salt Lake City, UT) amplifier from either one or both FFAs of each participating subject, using a Blackrock Microsystems Cabrio headstage to current-amplify and re-reference each channel to the reference wire in its respective bundle. Recordings took place between one and three days after implant to minimize the possibility of microwire damage and unit dropout.

### Tasks

All subjects performed the 1-back task shown in Fig. S2. Sets of ten faces, body parts, houses, patterns and tools were presented centrally, on a laptop monitor using Presentation software (Version 0.70, Neurobehavioral Systems, Berkeley, CA; picture size: ~12° at ~60 cm viewing distance), while subjects were sitting up in bed. Images were shown in blocks of a single category at a time. All ten exemplars from the relevant category were presented once in each block in a pseudo-random order, with the exception of 1-back repeats where the same exemplar was presented twice. Each image was presented for 0.50 s, followed by a jittered inter-stimulus period of between 0.75 and 1.5 s. Blocks were separated by either 4 or 8 s. Two versions of the task were used. One contained 250 trials (25 blocks), and was only performed by Subject 4, and the other contained 260 trials (26 blocks), and was performed by all other subjects. Each exemplar was presented 3–7 times throughout the task. There were 18–19 1-back repetitions during the task, for which the subjects were instructed to click a mouse button on the laptop on which the images were presented. Faces images were drawn from the dataset compiled by Minear and Park^36^.

The following are adapted from Norman, *et al*.^11^: four subjects (Subjects 2, 3, 6 and 8) performed the free recall task. The experiment was divided into two runs. Participants were presented with 14 different pictures of famous faces and popular landmarks. Each picture repeated four times (1.5 s duration, 0.75 s inter-stimulus interval) in a pseudorandom order, such that each presentation cycle contained all of the different pictures but the order of pictures was randomized within the cycle. The same picture was never presented twice consecutively. Participants were instructed to look carefully at the pictures and try to remember them in detail, emphasizing unique colors, unusual face expressions, perspective, and so on. Stimuli were presented on a standard laptop LCD screen using the Presentation software (Version 0.70, Neurobehavioral Systems, Inc., Berkeley, CA; picture size: 17° × 13° at ~60 cm viewing distance).

After viewing the pictures, participants put a blindfold on and began a short interference task of counting back from 150 in steps of 5 for 40 s. Upon completion, recall instructions were presented, guiding the subjects to recall items from only one category at a time, starting with faces in the first run and with places in the second run, and to verbally describe each picture they recall, as soon as it comes to mind, with two to three prominent visual features. The instructions also emphasized reporting everything that came to mind during the free recall period. The duration of the free recall phase was 2.5 min per each category (5 min in total, × two runs). In case the subjects indicated that they could not remember any more items, they received a standard prompt from the experimenter (*e.g*., “Can you remember any more pictures?”). The order of the recalled categories was fixed across subjects and counter-balanced between the two runs. A different set of pictures (7 per category) was presented in each run.

Verbal responses during the free recall phase were continuously recorded *via* a microphone attached to the subject’s gown. The onset and offset of each recall event were extracted in an offline analysis, identifying the first/last soundwave amplitude change relevant to each utterance using Audacity recording and editing software (version 2.0.6, https://www.audacityteam.org/).

### Spike Detection and Sorting

Combinato^37^ was used for spike detection and sorting in each of the channels. Candidate units were visually inspected for waveform consistency and plausible inter-spike interval distribution. When ambiguous, we preferred not to overscreen at this stage and to allow for subsequent objective analyses to reject artifacts.

### Identification of Visually-Responsive Units in the 1-Back Task

Spike trains were aligned to stimulus onset. The ON period was defined as 0.1 to 0.5 s after each stimulus appeared on screen and the OFF period was defined as 0.6 to 1 s after onset (0.1 to 0.5 s after the reappearance of the fixation cross). If at least one spike was not recorded during nine or more ON or OFF periods, then the unit was discarded from further analysis.

A Friedman’s test was used to determine whether there was a significant effect of time relative to stimulus onset on unit firing rates. The test was performed on the *n* by *i* array: |Δ*R_i_*[*n*]|, where *R_i_*[*n*] is the firing rate in each bin index *n* of width 0.033 s from 0 to 0.5 s after stimulus change (onset or offset), Δ is the first discrete differential, and *i* is the run number, used as the multiple measures factor in the Friedman’s test. The test is performed relative to onset and offset, and compared to α = 0.05 with Bonferroni correction for multiple comparisons across units. It has the effect of looking for changes in time-varying firing rates while ignoring amplitude differences across stimuli.

### Identification of Category-Selective Units in the 1-Back Task

A unit was defined as category selective if spike rates were significantly unequal among stimulus categories (sample sizes for Subjects 1–3 and 5–8: 60 face presentation trials and 50 presentations trials of all other stimulus categories; sample sizes for Subject 4: 50 presentation trials of all stimulus categories) during the ON and OFF periods for ON and OFF cells, respectively, according to a Kruskal-Wallis one-way analysis of variance test at α = 0.05 corrected for multiple comparisons across visually-responsive units using the Bonferroni-Holm technique. Firing rates among categories were further compared to establish if any one category showed significantly higher or lower firing rates than all others at α = 0.05, using the Fisher’s least significant difference procedure to account for multiple comparisons; if one category of stimuli evoked significantly stronger or weaker responses, then the unit was said to be selective for that category of stimuli, and “excited” or “inhibited”, respectively. If a unit simultaneously showed significant increases and decreases to different categories, then the magnitudes of the responses (absolute values of the natural-log-ratios of the spikes rates during the ON period and the 0.4 s before stimulus onset for ON cells and during the ON and OFF periods for OFF cells, with 0.1 Hz pseudocounts added before taking the log) is compared using a Wilcoxon rank-sum test with stimulus categories as groups. If responses to one category are significantly stronger than to the other at α = 0.05 (two-tailed), then the unit is said to be selective for that category.

### Calculation of Unit Response Latency

Response latency of face-selective units was calculated by constructing peristimulus-time histograms with bin widths of 0.02 s for each face-selective unit, and comparing the activity 0 and 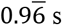 post-onset to *Z*^−1^(0.95) × *S*_BL_ for ON cells and *Z*^−1^(0.05) × *S*_BL_ for OFF cells, where *Z*^−1^ is the inverse normal cumulative distribution function, and *S*_BL_ is the sample standard deviation of the firing rate among the bins between 0.5 and 0 s before stimulus onset. The center of the first of three consecutive bins exceeding this threshold is defined as the onset latency for that unit.

### Relationship Between Units’ Visual Responsiveness and Category Selectivity

The *χ*^2^ parameter from the Friedman test used to measure unit responsiveness (responsiveness index) is plotted against the *χ*^2^ parameter from the Kruskal-Wallis test used to measure unit category selectivity (selectivity index) on a log-log scale. We used only subjects with the same distribution of trials per exemplar so that category selectivity values could be directly compared. The Spearman correlation between the log values of the two indices is calculated and tested against zero at α = 0.05.

The procedure is repeated using the *d*’ face sensitivity index instead of the *χ*^2^ Kruskal-Wallis value.

### Exemplar Decoding

Log-firing rates (pseudocount 1 added before taking the log) from pseudo-populations of visually-responsive units from all eight subjects, at 0.1 to 0.5 s after image presentation were calculated for three random trials of two exemplars. Two of the three trials from each exemplar are used as training data, and the third as a test probe (this is done because the least-repeated exemplar has three trials). The pseudo-population response vectors are transformed into 2-dimensional space using the classical multidimensional scaling algorithm in MATLAB (version 2018a, `cmdscale()`), which linearly transforms the data into a two-dimensional space while optimally preserving Euclidean distance relationships between trials. A Fisher’s linear discriminant is fit to the training data, and the held-out test trials (one from each exemplar) are classified, resulting in an accuracy per iteration of 0, 0.5 or 1.0. This process was repeated for 10000 iterations for each pair of exemplars, and the mean accuracy for each pair of categories, and within each category, for each iteration are compared to a bootstrap distribution of 10000 mean accuracies obtained by performing the same classification procedure on data with labels shuffled among the trials within the category (for within-category exemplar decoding) or among the categories (cross-category exemplar decoding) being compared. If the median rank exceeds 95% of the bootstrap distribution, Bonferroni corrected at *N* = 15 (for 5 within-category and 10 cross-category comparisons), the exemplars within the category or between categories are decoded above chance.

### Multidimensional Scaling Plots

Log-firing rates from pseudo-populations of all eight subjects in response to the first three house and face exemplars, normalized (*Z*-scored) within each unit among trials, were transformed using the classical multidimensional scaling algorithm in MATLAB (`cmdscale ()`) to 2-dimensional representations, first exemplars of both categories together, and then of houses and faces, alone.

### Free Recall Preprocessing

Spike trains were aligned to stimulus onset. Firing rates for each candidate unit during key intervals (−0.1 to 0.1 s, 0.1 to 0.3 s, 0.3 to 0.5 s, and 0.5 to 1.6 s relative to image presentation) for face and place images were passed as input to a Friedman’s test with trial number as the repeated measures factor. If *p* < 0.025 (Bonferroni corrected at *N*=2 for face and place categories) the unit was classified as visually-responsive by the standard criterion, and if *p* < 0.001, it was classified as visually-responsive by the strict criterion. A Wilcoxon rank-sum test was used to identify differences in firing rate between image categories on visually-responsive unit spike trains 0.1 to 0.5 s post-stimulus-onset at α = 0.025, and units with significant differences are classified as category selective.

Category preferences (face sensitivity, *d*’) of units were calculated as follows:

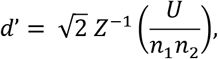

where *Z*^−1^ is the inverse normal cumulative distribution function, *U* is the Mann-Whitney *U* statistic, and *n*_1_ and *n*_2_ are the number of place and face presentation trials, respectively. Face-selective units are redefined as those with a *d*’ greater than that of the greatest *d*’ associated with a non-category-selective unit by the rank-sum test (as above).

### Free Recall Calculation of Units’ Category Preference

Spike trains were aligned to free recall speech. Mean firing rates during the time period between −2 and 2 seconds relative to utterance onset were calculated for each utterance and each unit, and the face sensitivity of spike rates associated with face and place utterances were calculated as above.

### Selectivity of Visually-Responsive Units During Free Recall

The recall category preferences of units classified as visually-responsive, according to the standard criterion, were compared to their respective category selectivities during image presentation. If the Spearman correlation coefficient was different from 0 at α = 0.05, then the results were confirmed using only units that exceed the strict criterion for visual responsiveness.

For additional confirmation, the peri-recall time histogram (with bins of 0.1 s width) of each face-selective unit was divided by the mean firing rate of that unit over all recall periods. The baseline-corrected peri-recall time histograms for all face-selective units recorded for each recall event from each subject-implant were averaged together and convolved by a 1-s boxcar (the result is a mean response from face-selective units that is relatively independent of clustering decisions and the number of units in each subject). A two-tailed unpaired *t*-test was used to compare mean activity −2 to 2 sec relative to the onset of the set of face and place recollection utterances, respectively.

Repeated *t*-tests was used to compare adjusted firing rates, downsampled to 2 Hz, at α = 0.05, at each time point, without correction.

Changes in firing rate during face recall (−2 to 2 s relative to utterance onset compared to mean firing rate over face recall blocks) in each visually-responsive unit (standard criterion) were compared to the changes during face image presentation (firing rate 0.1 to 0.5 s minus −0.4 to 0 s relative to image onset). Spearman’s ρ and the parameters of the best fit line are calculated, and the correlation is tested against zero at α = 0.05.

### Relationship Between Firing Rates During Presentation and Recall

Mean firing rates for each visually-responsive (standard criterion) unit from each subject were calculated for −2 to 0 s relative to onset of face recall utterances. Mean firing rates from the same exemplars during presentation were weighted by the number of utterances in which they were referenced. Each visually-responsive unit is plotted on a log-log scale based on presentation and recall firing rates. A first-degree polynomial is fit to the log-transformed data (0.01 pseudocount added to avoid zeros), and *R*^2^ fit is calculated, with the associated *p* value compared to 0.05.

### Exemplar Decoding of Stimuli from Free Recall

The procedure for exemplar decoding in the 1-back task was repeated for data from the presentation phase of the free recall experiment, using mean firing rates between 0.1 and 0.5 s after image onset, to calculate decoding accuracy on an individual-subject level. Each exemplar was shown four times, so each classifier was trained on 3 trials of each exemplar and tested on one. Additionally, as shown in Fig. S13, mean firing rates may differ between image presentation and recall, so to normalize each trial, the mean of each log-firing-rate pseudo-population vector was subtracted from each component of that vector, and all elements then divided by one greater than the standard deviation (to avoid divide-by-zero errors). Category and cross-category accuracies were averaged across exemplar pairs and blocks at each iteration, and compared to a bootstrap distribution obtained by shuffling the labels, as describe for the 1-back task. The two runs performed by Subject 3 were coupled together as if they had occurred consecutively. Ten thousand cross-validation runs were performed, each time randomizing the test trial for each exemplar. The median percentile rank of the classifier accuracy for both categories and the cross-category result for each subject were compared to a statistical threshold of α = 0.05, corrected for multiple comparisons using the Bonferroni-Holm technique (*N* = 4 subjects × 3 category domains), respectively. Only subjects and categories with significant presentation exemplar decoding were tested for recall exemplar decoding.

A population vector was built from each recall event, taking the log-base-10 (pseudocount 1) of the mean firing rates of the neural ensemble from each subject, 2 to 0 seconds before the onset of the utterance. The population vector was normalized (as above) and classified with each of the binary classifiers trained to discriminate between each stimulus pair by presentation response (as above, but using all four trials for training). To calculate the cross-category classification performance, the full classifier array was applied to each recall datum; to calculate within-category classification performance, only the classifiers trained on pairs of faces or pairs of places were used on their respective stimuli.

The number of times that the classifiers from the array classified each input stimulus as belonging to a given exemplar is rank ordered. If a recall event is classified as belonging to the two different exemplars the same number of times, the ranking is repeated recursively using only the classifiers trained on those exemplars until the tie is broken or the number of classifications per exemplar stabilizes (in which case, the exemplars are assigned the mean of the two ranks). The ranks associated with the correct exemplar in all recall events for each stimulus category (faces and places, respectively) and all recall events together (cross-category) are added across blocks for each subject and compared to a bootstrap distribution obtained by randomly permuting the labels of the recall events within each stimulus category, and repeating the classifier analysis 10000 times (place and face) or 2000 times (cross-category). Bootstrap distribution ranks of the accuracies of all subjects with significant category decoding during presentation were combined using Fisher’s method of combined probabilities, corrected for multiple comparisons using the Bonferroni-Holm method (among subject-categories with significant presentation exemplar decoding on the same task), and tested for significance at α = 0.05.

To confirm successful recall exemplar decoding, exemplars were split into presentation response terciles for each face-selective unit, with the top and bottom terciles defined as the sets of exemplars evoking mean presentation responses above the 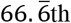 and below the 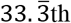 percentiles of presentation mean response magnitudes (firing rates 0.1 to 0.5 s after presentation) across all four presentation trials, respectively. Exemplars were then split into recall activity terciles, with the top and bottom terciles defined as the sets of exemplars preceded by mean recall activity above the 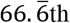 and below the 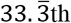 percentiles of mean recall activity magnitudes (mean firing rates 2 to 0 s before recall utterance) across all recall events associated with that exemplar. The mean, across face-selective units, of the fraction of exemplars in the top and bottom presentation response terciles that were also in the top and bottom recall activity terciles for each unit, respectively, relative to the total number of exemplars in the top and bottom recall activity terciles for that unit, was compared to a surrogate distribution obtained by pseudorandomly permuting the exemplar labels 10000 times. The faction of matching exemplars so obtained was compared to α = 0.05, with the Bonferroni-Holm correction for multiple comparisons (for the number subjects with face-selective units – 3). In this context, more matching exemplars is equivalent to more congruence between presentation and recall activity for any given exemplar. This process was then repeated with only face exemplars in those subjects in whom congruence was above chance. Subject 6 was excluded due to a lack of face-selective units.

## Acknowledgments

We thank Salman Qasim and Drs. Joshua Jacobs, Ida Momennejad, Elliot Smith, Ella Podvalny, Tal Golan, Pierre Megévand, José Herrero, Victor Du and Stephan Bickel for intellectual and technical support. We thank Dr. Xiangzhi Zhou for help with the fMRI experiments. We acknowledge the contributions of our colleagues at the Northwell Health Comprehensive Epilepsy Center, especially Willie Walker, R. EEG. T., Monika Lalik, R. EEG. T., and Drs. Sean Hwang, Fred Lado and Scott Stevens. We would like to thank our patient-subjects and their families, without whose patience and cooperation, this research would have been impossible.

## Funding

A.D.M. receives funding from the following grants: NIH/NINDS NS098976-01, NSF-BSF-2017015; NIMH MH114166-01; S.K. is supported by the Feinstein M.D. Scholarship; This work was partly funded by US-Israel BSF grant to A.D.M. and R.M.;

## Author contributions

All authors contributed to conceptualization of this research. S.K. performed formal analysis on electrophysiological data, and E.M.Y. and S.G. on imaging data. S.K. and E.M.Y. collected the data. S.K. and Y.N. developed the methodology. A.D.M. performed all of the surgeries. A.D.M. and R.M. provided resources and supervision. S.K. and E.M.Y. created visualizations. S.K. and A.D.M wrote the original draft; all authors reviewed and edited the manuscript.

## Competing interests

A.D.M. has a consulting agreement with Medtronic.

## Data and materials availability

Data will be provided upon request. Code is available on: https://github.com/IEEG/SUFreeRecall.

## Supplementary Materials

Supplementary Text

Figs. S1 to S12

Tables S1 and S2

## Notes

#### Summary of Updates

Text updated, new analysis included (concordance between unit exemplar tuning during presentation and recall).

https://github.com/IEEG/SUFreeRecall

