## Supplemental Materials for "Face-selective units in human ventral temporal cortex reactivate during free recall"

### Face-selective units in human FFA reactivate during face recall and imagery

**This PDF file includes:**

Supplementary Text
Figs. S1 to S12
Tables S1 and S2

**Other Supplementary Materials for this manuscript include the following:**

**Supplementary Text**

Behavioral performance on 1-back task

All subjects were attentive during the 1-back task. Subject 2 performed the task twice: once for each implant. Subjects missed between 5 and 67% of 1-back repeats (button not pressed within 1.5 s, median of all runs of all subjects: 47%). Between 5 and 69% (median: 47%) of button presses were false detections (not in the 1.5 s following a duplicate stimulus).

Relationship Between Responsiveness and Selectivity

Figure S4 shows that visual responsiveness is correlated with category selectivity, generally, and face selectivity, specifically ($p < 0.001$, Spearman's ρ, both). The distribution shows that while there are some strongly-responsive units with poor face selectivity, there are no face-selective units that show poor responsiveness. It is unsurprising that units that responded strongly to images of any kind are more likely to show a contrast among the different categories if category selectivity exists. Similarly, if a preference for faces exists among the population, it would emerge as a positive correlation between visual responsiveness and face *d*' sensitivity. Together, these results reinforce our claim that category preferences exist in the ventral temporal cortex, and in our population of FFA-targeting electrodes, face-selectivity dominates. This result also gives us an additional means by which to double-check our results by testing them on a smaller set of units passing a stricter threshold for visual responsiveness (as in Fig. 3B and C).

Free Recall Behavior

Of the four subjects, only Subject 2 showed poor performance on the first run of the free recall task. Upon inquiry, it was established that this subject did not comprehend the instructions. The instructions were repeated, and the second run showed satisfactory performance. Subject 2 was also implanted with a second electrode in the contralateral FFA, and the first run of the task was repeated with different stimuli. The two individual runs are analyzed together for the exemplar decoding portion of the analysis because two data sets obtained from contralateral hemispheres of the same subject are not necessarily expected to be independent.

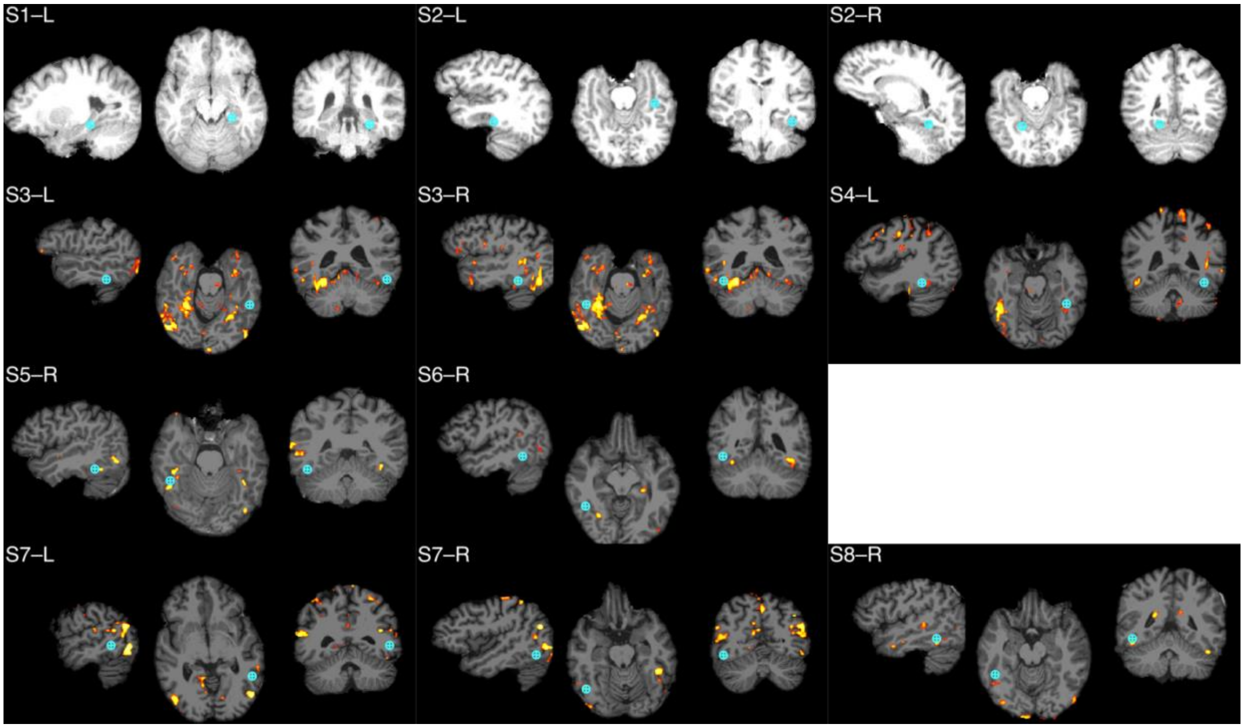


**Fig. S1.**

Electrode localizations from all subjects. Subjects 1–8 labeled S1–8, with L and R indicating left and right electrodes respectively. Crosshairs centered at terminal macro contact, with surrounding circles representing the 4 mm radius of the potential microelectrode range. Where functional imaging is available (Subject 3–8), yellow/red areas represent regions with face>house responses, except in Subject 4, in which face>pattern responses were used.

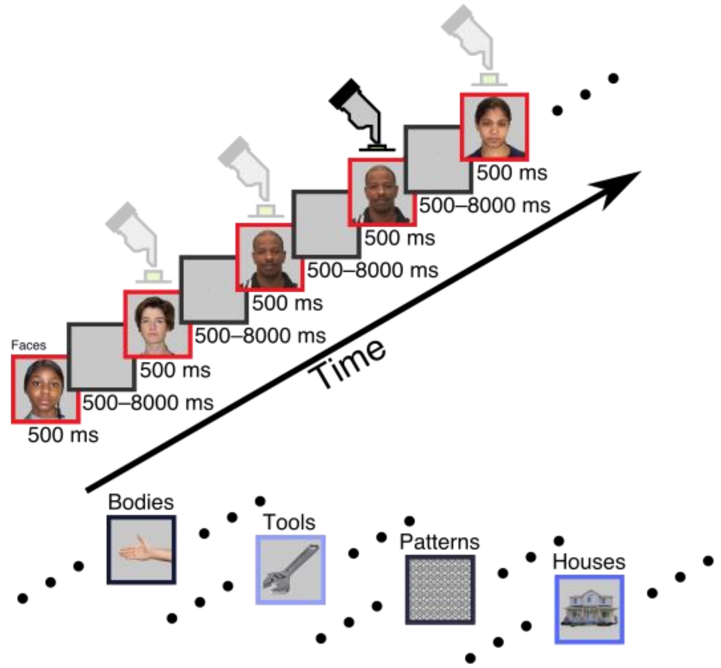


**Fig. S2.**

One-back task design. Subjects were shown a series of images of one of five categories: faces, bodies, houses, patterns and tools. Each image was displayed for 500 ms. There was a random inter-stimulus interval of 500 to 8000 ms. Blocks of images of the same category were shown together. Subjects were instructed to press the button when they saw the same image twice in a row.

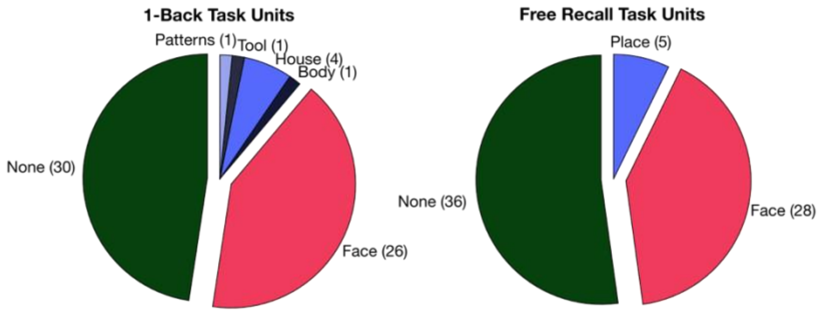


**Fig. S3.**

Pie chart showing the distributions of category selectivity among visually-responsive units from all participating subjects in the 1-back and free recall/imagery tasks. The overwhelming majority of units that were selective for a single category were selective for faces.

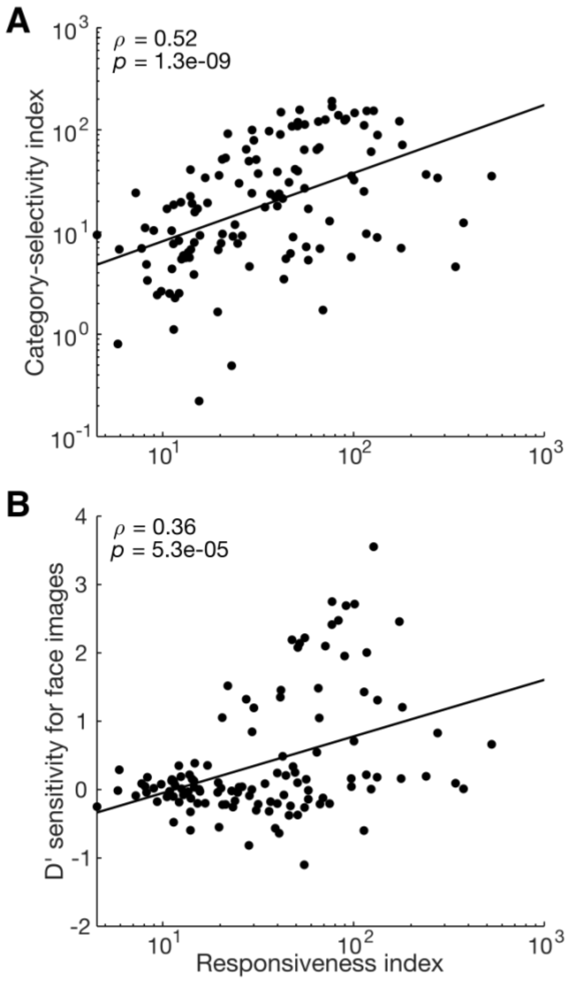


**Fig. S4.**

Correlation between unit visual responsiveness and (**A**) category selectivity and (**B**) *d*' face sensitivity over other categories. Visual responsiveness and category preference are strongly correlated ($p < 0.001$, Spearman's ρ, $N = 120$ units in subjects who performed the standard version of the task).

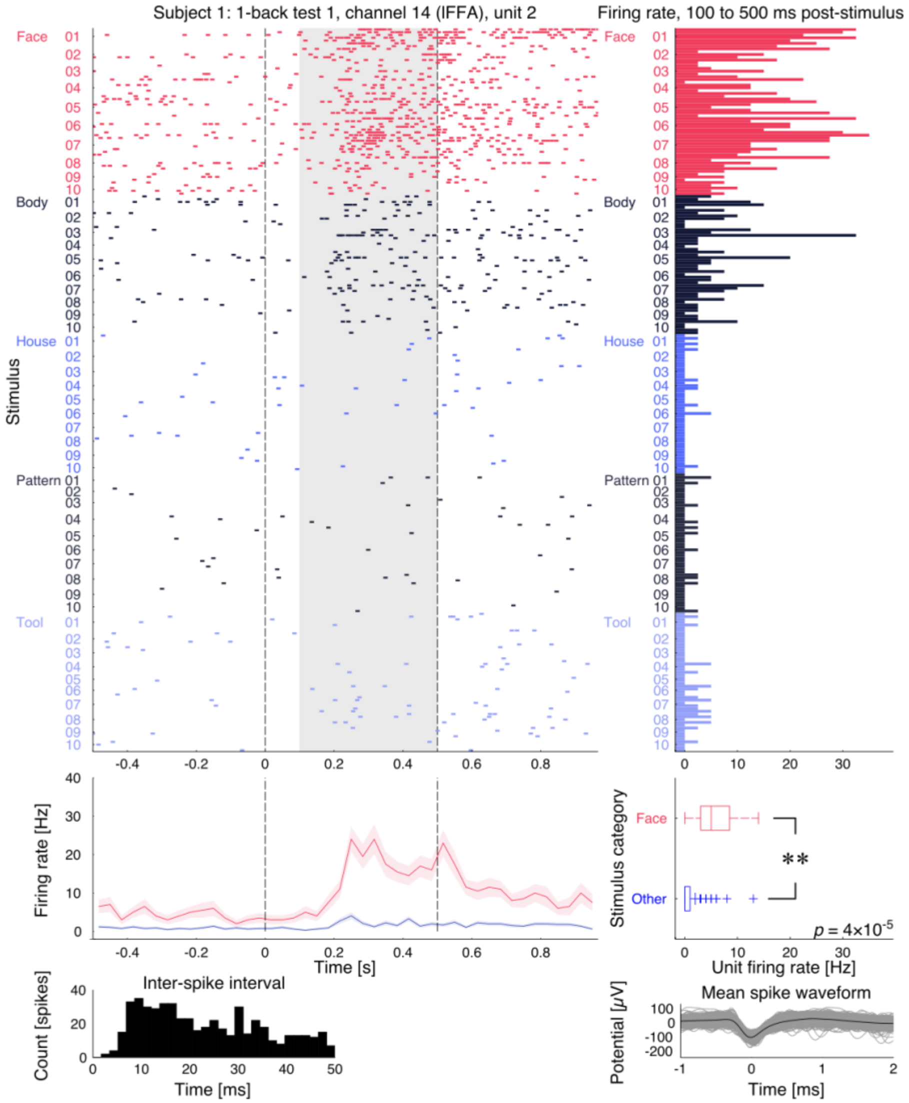


**Fig. S5.**

Sustained response to faces. Expanded data from Fig. 1E, responses of Subject 1, channel 14, unit 2 (lFFA) to image presentation during the 1-back task. Raster plot: each row represents a trial, vertical dashed lines represent image onset (0 seconds) and offset (0.5 s). The mean firing rate for each trial over the gray shaded area (0.1 to 0.5 s) is shown in the bar graph at right, with a box plot summarizing firing rates of all trials directly below. Firing rates following face image presentation were significantly higher than those following non-face object images (Wilcoxon rank-sum test, $p < 0.001$). Peristimulus time histogram shown in line graph below raster plot, with colored areas representing mean ± standard error of the mean for each timepoint (not corrected for multiple comparisons), showing selectively-elevated firing rate in response to face images (red) relative to non-face object images (blue) starting approximately 0.1 s after presentation and remaining elevated above baseline after image offset. On the bottom are a histogram of inter-spike intervals (left) and superimposed spike waveforms (gray) and mean spike waveform (black) for this unit (right), both showing plausible single-unit characteristics. **: $p < 0.001$. lFFA: left fusiform face area.

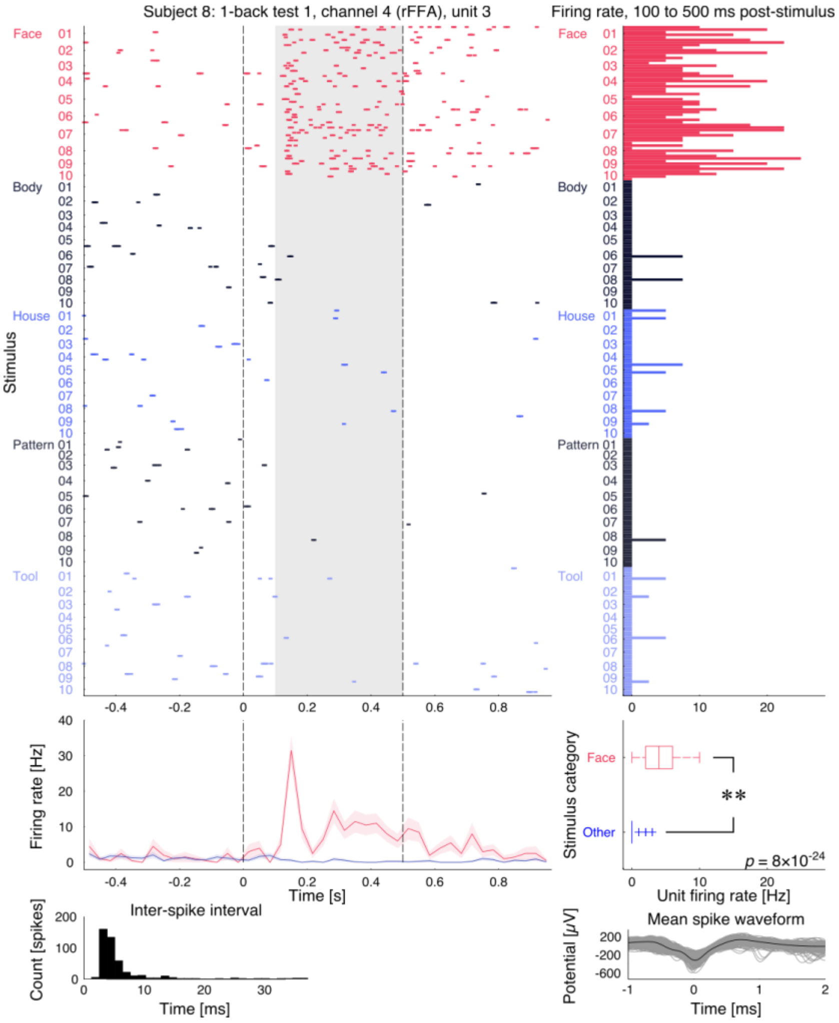


**Fig. S6.**

Transient peak in activity to face presentation. Expanded data from Fig. 1F, responses of Subject 8, channel 4, unit 3 (rFFA) to image presentation during the 1-back task. Firing rate dips and resumes after peak at a lower, sustained rate. **: $p < 0.001$, rank-sum test. rFFA: right fusiform face area.

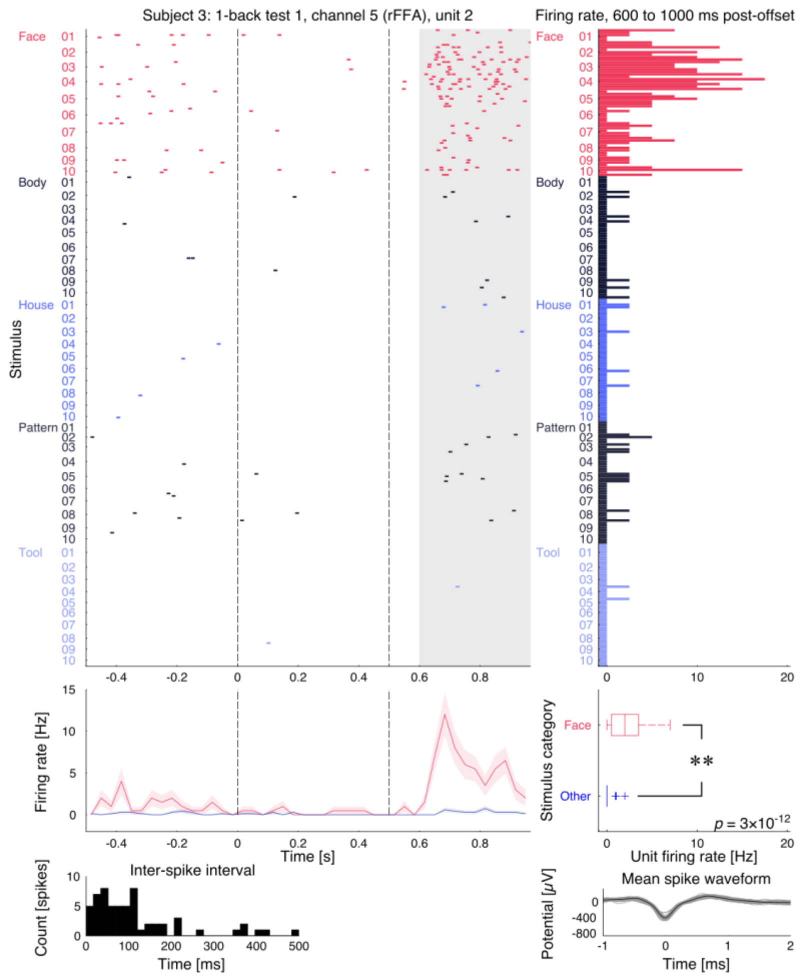


**Fig. S7.**

Offset response to faces. Expanded data from Fig. 1G, responses of Subject 3, channel 5, unit 2 (rFFA) to image presentation during the 1-back task. The mean firing rate for each trial over the gray shaded area (0.6 to 1 s) is shown in the bar graph at right of the raster, with a box plot summarizing firing rates of all trials directly below. **: $p < 0.001$, rank-sum test. rFFA: right fusiform face area.

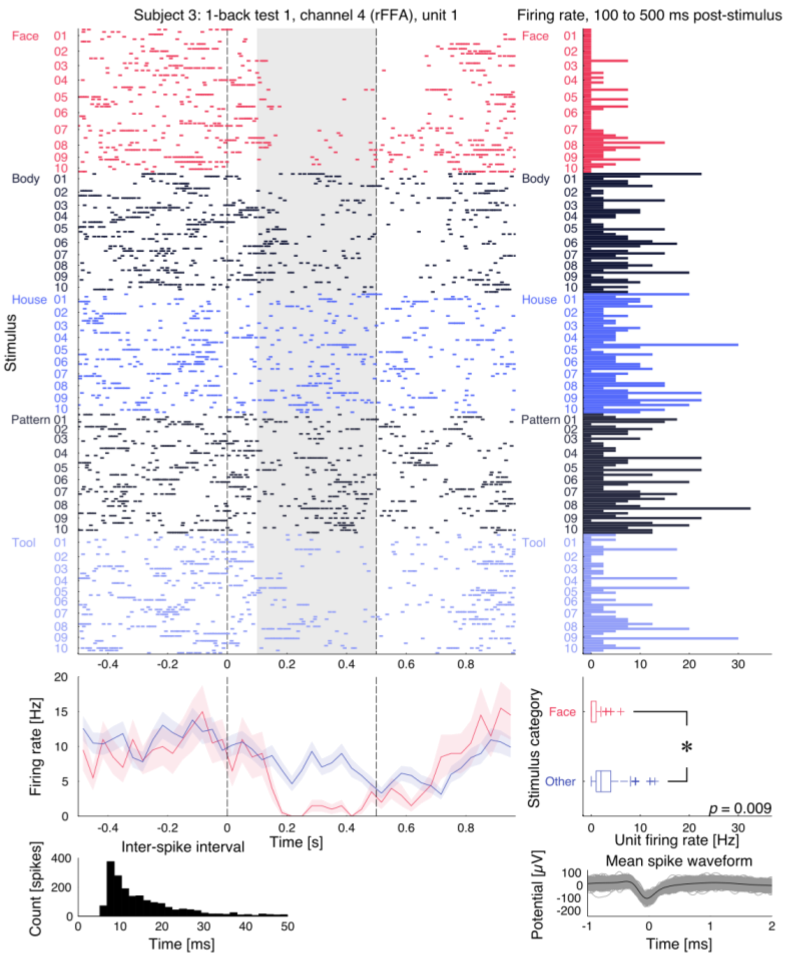


**Fig. S8.**

Suppression to faces. Expanded data from Fig. 1H, responses of Subject 3, channel 4, unit 1 (rFFA) to image presentation during the 1-back task. This unit also demonstrates bursting behavior. *: $p < 0.05$, rank-sum test. rFFA: right fusiform face area.

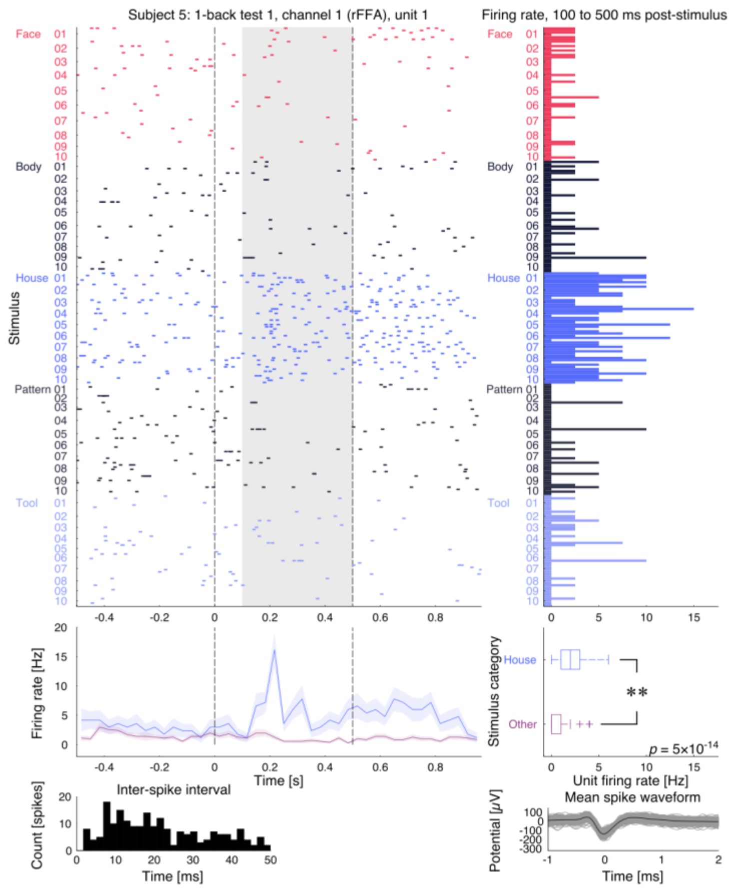


**Fig. S9.**

House-selective unit. Expanded data from Fig. 1I, responses of Subject 5, channel 1, unit 1 (rFFA) to image presentation during the 1-back task, demonstrating a house-selective unit. Peristimulus time histogram below raster plot shows elevated firing rates in response to house images (blue) relative to all other (purple). Boxplot confirms greater firing rates between 0.1 and 0.5 s after stimulus onset. **: $p < 0.001$, rank-sum test. rFFA: right fusiform face area.

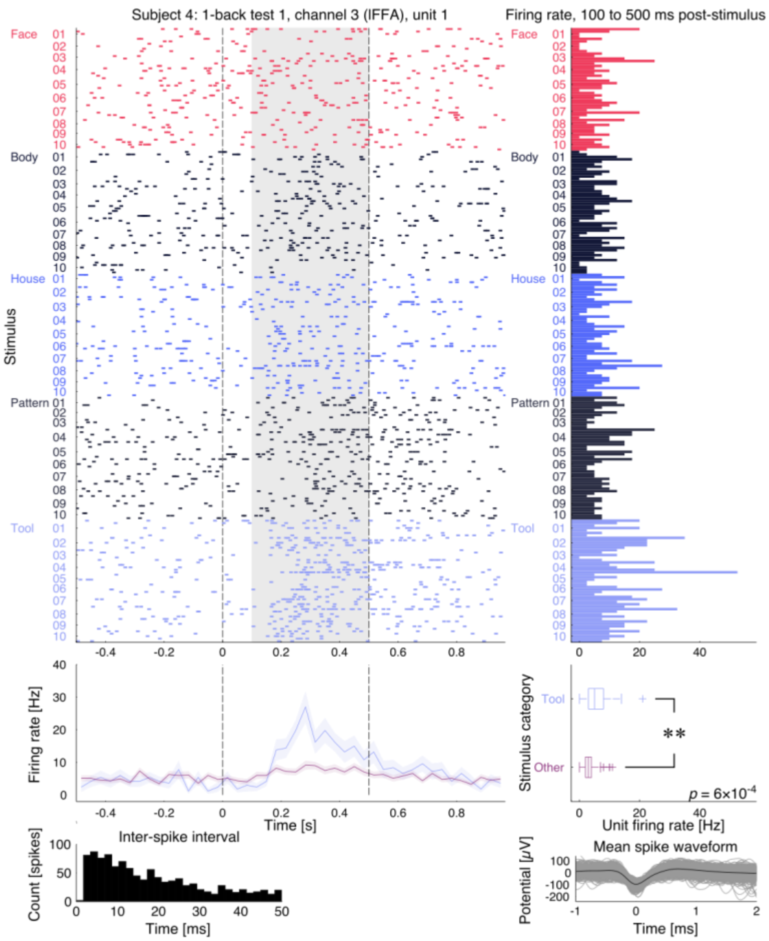


**Fig. S10.**

Tool-selective unit. Expanded data from Fig. 1J, responses of Subject 4, channel 3, unit 1 (lFFA) to image presentation during the 1-back task. Peristimulus time histogram below raster plot shows elevated firing rates in response to tool images (blue) relative to all other (purple). Boxplot confirms greater firing rates between 0.1 and 0.5 s after stimulus onset. **: $p < 0.001$, rank-sum test. lFFA: left fusiform face area.

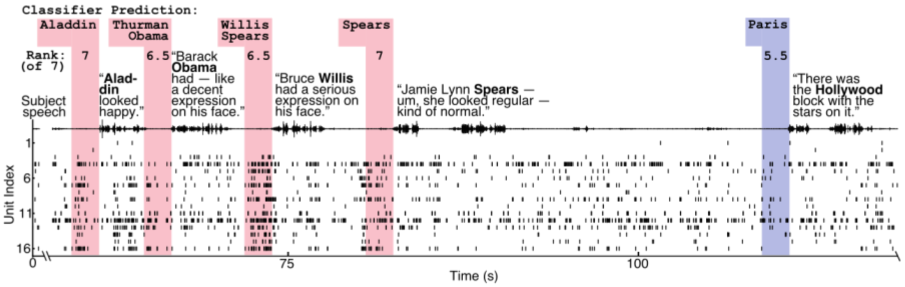


**Fig. S11.**

Audio and neural spike raster plot from an example face recall run from Subject 8. Excerpt shows four face recall events (red) and one place recall event (blue), with speech audio envelope waveform and raster plot of 16 visually-responsive and face-selective units. Colored boxes highlight the 2 s prior to onset of each recall utterance, unit firing rates during which serve as the input for the classifier, and show the top classifier prediction (or top two predications if tied for first), and, below, the rank of the correct exemplar when voted upon by the full set of classifiers, out of 7, where 7 is the best performance. Excerpts of subject speech with face or place identity in bold. The four face exemplars recalled by the subject rank either as the top classifier-set prediction, or are tied for first. The place exemplar is not predicted.

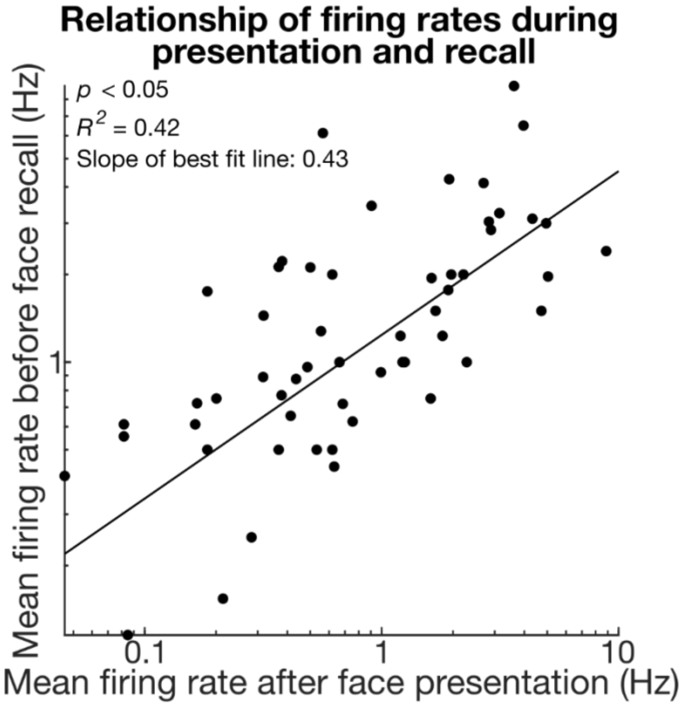


**Fig. S12.**

Mean firing rates during recall do not necessarily match those during presentation. Weighted mean of firing rates during presentation (0.1 to 0.5 s after onset) of remembered faces is plotted against mean firing rate before recall (-2 to 0 s relative to start of utterance) for visually responsive units ($N = 62$). For units that show strong activity during face presentation, average activity during face recall appears to be modestly weaker. This justifies using normalized firing rates as input to the classifiers trained on free recall and imagery task data. It does not imply that peak instantaneous firing rates are greater during presentation than recall. The slope is significantly different from zero ($p < 0.05$), as expected.

| Subject | 8 | 2 | 3 | 6 |
|---|---|---|---|---|
| Faces | 0.006 | 0.019 | <0.001 | 0.12 |
| Places | 0.3 | 0.019 | 0.5 | 0.3 |
| Inter-Category | <0.001 | <0.001 | <0.001 | 0.09 |

**Table S1.**
*P* values associated with Fig 4B, in the main text.

| Subject | 8 | 2 | 3 | 6 |
|---|---|---|---|---|
| Faces | 0.018 | 0.6 | 0.04 | 0.9 |
| Places | 0.07 | 0.17 | 0.18 | 0.4 |
| Inter-Category | <0.005 | 0.5 | 0.009 | 0.6 |

**Table S2.**

*P* values associated with Fig 4C, in the main text.
